# Bacterial, phytoplankton, and viral dynamics of meromictic Lake Cadagno offer insights into the Proterozoic ocean microbial loop

**DOI:** 10.1101/2021.10.13.464336

**Authors:** Jaspreet S Saini, Christel Hassler, Rachel Cable, Marion Fourquez, Francesco Danza, Samuele Roman, Mauro Tonolla, Nicola Storelli, Stéphan Jacquet, Evgeny M. Zdobnov, Melissa B. Duhaime

## Abstract

Lake Cadagno, a permanently stratified high-alpine lake with a persistent microbial bloom in its anoxic chemocline, has long been considered a model for the low-oxygen, high-sulfide Proterozoic ocean where early microbial life gave rise to Earth’s oxygenated atmosphere. Although the lake has been studied for over 25 years, the absence of concerted study of the bacteria, phytoplankton, and viruses, together with primary and secondary production, has hindered a comprehensive understanding of its microbial food web. Here, the identities, abundances, and productivity of microbes were evaluated in the context of Lake Cadagno biogeochemistry. Photo-synthetic pigments and chloroplast 16S rRNA gene phylogenies suggested high abundances of eukaryotic phytoplankton, primarily *Chlorophyta*, through the water column. Of these, a close relative of *Ankyra judayi*, a high-alpine adapted chlorophyte, peaked with oxygen in the mixolimnion, while *Closteriopsis*-related chlorophytes peaked in the chemocline and monimolimnion. Anoxygenic phototrophic sulfur bacteria, Chromatium, dominated the chemocline along with *Lentimicrobium*, a newly observed genus of known fermenters. Secondary production peaked in the chemocline, suggesting anoxygenic primary producers depended on heterotrophic nutrient remineralization. Virus-to-microbe ratios spanned an order of magnitude, peaking with high phytoplankton abundances and at a minimum at the peak of Chromatium, dynamic trends that suggest viruses may play a role in the modulation of oxygenic and anoxygenic photo- and chemosynthesis in Lake Cadagno. Through the combined analysis of bacterial, eukaryotic, viral, and biogeochemical dynamics of Lake Cadagno, this study provides a new perspective on the biological and geochemical connections that comprised the food webs of the Proterozoic ocean.

**IMPORTANCE:** As a window to the past, the study offers insights into the role of microbial guilds of Proterozoic ocean chemoclines in the production and recycling of organic matter of sulfur- and ammonia-containing ancient oceans. The new observations described here suggest that eukaryotic algae were persistent in the low oxygen upper-chemocline in association with purple and green sulfur bacteria in the lower half of the chemocline. Further, this study provides the first insights into Lake Cadagno viral ecology. High viral abundances suggested viruses may be essential components of the chemocline where their activity may result in the release and recycling of organic matter. The framework developed in this study through the integration of diverse geochemical and biological data types lays the foundation for future studies to quantitatively resolve the processes performed by discrete populations comprising the microbial loop in this early anoxic ocean analogue.

## INTRODUCTION

Roughly 2.3 billion years (Gyr) ago the oceans were filled with sulfur (1) and anoxygenic photosynthesis was a dominant mode of microbial primary production (2). Following the great oxidation event 2 Gyr, anoxygenic photosynthesis continued to sustain microbial life in the euxinic (anoxic, high sulfur) environments below oxygenated surface waters (1, 2, 3). Habitats in which anoxygenic photosynthesis significantly contributes to primary production continue to persist, such as in anoxic zones of permanently stratified sulfur-rich meromictic lakes, like Lake Cadagno (4) where hydrogen sulfide serves as an important electron donor (5). These permanently stratified lakes are resistant to seasonal mixing events that would otherwise redistribute oxygen throughout the water column (6). In these systems, biogeochemical processes provide a link between the contrasting dominant modes of primary production, thereby directly connecting the food webs of the oxygenated and anoxic habitats.

The microbial community composition of meromictic Lake Cadagno typifies the reconstructed signatures of life in the Proterozoic ocean (7, 8). Purple and green sulfur bacteria (PSB and GSB, respectively) are amongst the hallmark bloom-forming microbes that are frequently observed in the chemocline of Lake Cadagno (9, 10). Within the chemocline, carbon, nitrogen, sulfur, and iron cycling are integrated through metabolic activities of the anoxygenic photosynthetic PSB and GSB (11, 12, 13, 14). Two PSB, *Thiodictyon syntrophicum* and *Chromatium okenii*, have been identified as the greatest contributors to primary production in Lake Cadagno (10, 15, 14). However, recent measurements of primary production to date have been limited to the chemocline (16, 10) and measurements of secondary production do not exist. Phytoplankton (including modern eukaryotes and cyanobacteria) likely coexisted with anoxygenic phototrophic sulfur bacteria in Proterozoic oceans (17, 18). High phytoplankton abundances and productivity were expected in the oxic zone because phytoplankton produces organic matter via oxygenic photosynthesis. In order to better understand microbial loop linkages across the stratified mixolimnion-chemocline transition in Lake Cadagno, paired measurements of productivity, microbial community composition, and diversity are essential next steps.

Lake Cadagno studies to date have focused on the phytoplankton and prokaryotic members of the microbial community and have yet to include viruses. Viruses are known to impact both phytoplankton and heterotrophic microbial populations through lysis and lysogeny and may hold a central position in the marine food webs and biogeochemical cycling (19, 20). Through the lysis of their microbial hosts, viruses play a role in the conversion of particulate organic carbon to dissolved organic carbon, which is predicted to be capable of sustaining the microbial loop (21, 22). Viruses have been identified in meromictic lakes (23, 24, 25), but their abundances across the vertical water column of Lake Cadagno are unknown.

To better understand the role the microbial loop plays in sustaining the food web across the stratified layers of Lake Cadagno, microbial community members (phytoplankton, prokaryotes, and viruses) were studied in the context of biogeochemical and primary and secondary production measurements across the oxic-anoxic transitions of the meromictic Lake Cadagno. We hypothesized the mixolimnion to have low primary production relative to the chemocline because the chemocline is known to be inhabited by a persistent microbial bloom of primary producing phototrophic sulfur bacteria (10). We also expected high secondary production and viral abundance to be an indicator of effective organic matter recycling within the Lake Cadagno water column, because photoautotrophs, both phototrophic sulfur bacteria and phytoplankton, rely on heterotrophs and viruses for the remineralization of organic matter and nutrient cycling (26, 27). Through a combination of physical, chemical and biological analyses, this work provides new evidence on how transitions between permanently stratified lake habitats and assemblages may sustain microbial food webs, informing our understanding of the aquatic ecosystems of early Earth.

## RESULTS

### Position of Mixolimnion, Chemocline, and Monimolimnion

Transitions between the major stratified layers sampled from Lake Cadagno were visually apparent by the colour of pigmented biomass captured on the 0.22 μm filters (Figure 1A). From the surface to 12.5 m depth, the oxygen levels fell from a maximum of 9.24 mg/L to 2.79 mg/L (light green, day 2; Figure 1B). As hypoxia is defined as <2 mg O_2_/L (28), 12.5 m marks the boundary of the oxic mixolimnion zone and the upper chemocline (top light purple; Figure 1). The chemocline was 13.5-15.5 m, as determined based on near-zero oxygen and light levels (day 2), peaked turbidity, a slight decrease in temperature, and a simultaneous rise in conductivity (dark purple; Figure 1B-D). At 14.5 m, oxygen levels were below detection and <1% of surface light penetrated (day 2; Figure 1B, D). Within the chemocline, the deep chlorophyll maximum (DCM) and peak turbidity shifted overnight between days 1 and 2, but in both cases, the peak turbidity was observed 0.5-1.5 m below the DCM (day 2; Figure 1B, D). Peak phycocyanin concentrations (26.53 μg/L) and basal fluorescence (Fb = 66.0) were observed at the DCM (day 2, 13.5-14 m) and above a secondary chlorophyll peak at 15 m (Figure 1D, Fig. S2). The monimolimnion was defined as the zone between 15.5 m and the benthos. The peak in particulate sulfur (0.74 ppm), H_2_S (2.84 mg/L) and ammonia (262.4 μg/L) concentrations were observed in the monimolimnion, as well as a decline in turbidity and photosynthetic pigments (Figure 1B, E, H).

**FIG 1.**
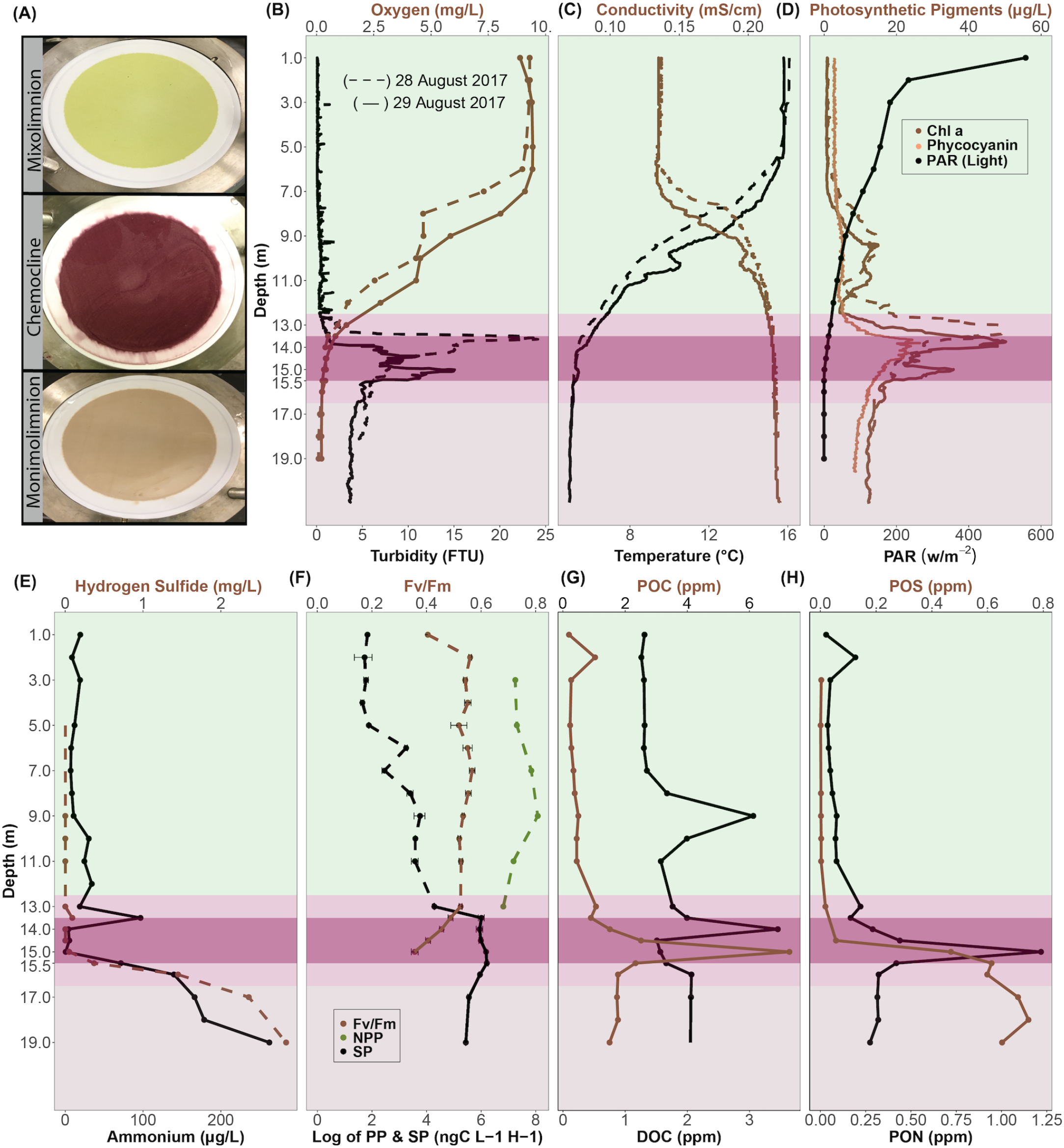
Depth profiles of biogeochemical parameters of Lake Cadagno. (A) Photographs of biomass collected on 0.22 μm (142 mm diameter) filters from Lake Cadagno mixolimnion, chemocline, and monimolimnion strata. The depth profiles of (B) oxygen and turbidity, (C) conductivity and temperature, (D) Chl a, phycocyanin, and photosynthetically active radiation (PAR) (E) hydrogen sulfide and ammonium concentrations, (F) net primary production (NPP) and secondary production (SP) rates, (G) particulate organic carbon (POC) and dissolved organic carbon (DOC) concentrations, (F) particulate organic sulfur (POS) and particulate organic nitrogen (PON) concentrations. In B-H, the mixolimnion, chemocline transition zone, chemocline, and monimolimnion layers are indicated by light green, lilac, purple and light brown backgrounds, respectively. Dashed and solid lines represent 28 and 29 August 2017, respectively.

### Microbial production at the oxic/anoxic interface

Between 7.0-11.0 m in the lower mixolimnion, there was a slight rise (up to 13.12 μg/L on day 1 and 11.80 μg/L on day 2) in Chl a (Figure 1D). The Chl a peak in the mixolimnion at 9.0 m corresponded with a peak in net primary production (NPP; 3210 ng C L-1 h-1; Figure 1D, F). The NPP peak declined in the lower mixolimnion (11.0 m) and upper chemocline (13.0 m). Though NPP was not measured below 13.0 m, basal and maximum fluorescence measured through the chemocline gave an indication of phytoplankton photosynthetic activity and efficiency (Fig. S2). In the oxic-anoxic boundary, a maximum Fv/Fm (an indicator of photosynthetic efficiency) of 0.49 was observed between 13.0-13.5 m and declined to 0.36 at 15.0 m (Figure 1F) where the light was absent. Overlapping with this oxygenic photosynthesis, the highest rates of secondary productivity (SP) were observed through the chemocline (Figure 1F). SP was significantly positively associated with some indicators of both photosynthetic microbes (Pearson: Chl a: R = 0.96, p-value = 1.7e-5; phycocyanin: R = 0.80, p-value = 0.005) and total biomass (Pearson: turbidity: R = 0.78, p-value = 0.007; total cell counts: R = 0.65, p-value = 0.04) (Fig. S3). In the chemocline, SP and indicators of NPP (phycocyanin, Chl a) were significantly negatively correlated with oxygen and light (Pearson: R = −0.7 to −0.9, p-values = 0.0002 to 0.01; Fig. S3). The peak of DOC (14.0 m) was followed by peaks of POC, PON (15.0 m; Figure 1G-H). In the monimolimnion below the chemocline, some organic and inorganic compounds continued to rise (POS, H_2_S, NH_4_^+^), while PON and POC dropped (Figure 1).

### Prokaryote- and Virus-like particle abundances and trends

The concentration of prokaryotic-like particles (PLPs; prokaryotic cell counts inferred from flow cytometry data) was highest in the chemocline with a peak of 905,000 cells/ml ± 28,583 at 15.0 m (496,000 ± 17,295 avg cells/ml), roughly half that in the monimolimnion (435,833 ± 10,243 avg cells/ml), and was lowest in the mixolimnion (109,556 ± 3,496 avg cells/ml). In contrast, virus-like particles (VLPs; viral counts inferred from flow cytometry data) were highest at the bottom of the mixolimnion (1.71e+08 ± 2.63e+07 VLP/ml at 11.0 m), lowest in the upper mixolimnion (7.03e+07 ± 1.50e+07 VLP/ml at 5.0 m), and relatively invariable through the rest of the water column (Figure 2B). As a result, the virus to microbe ratio (VMR) peaked with viruses at the bottom of the mixolimnion (VMR of 1642 at 11 m), but sharply decreased (VMR of 158 at 15 m) when prokaryotic-like particles rose in the chemocline (Figure 2C).

**FIG 2.**
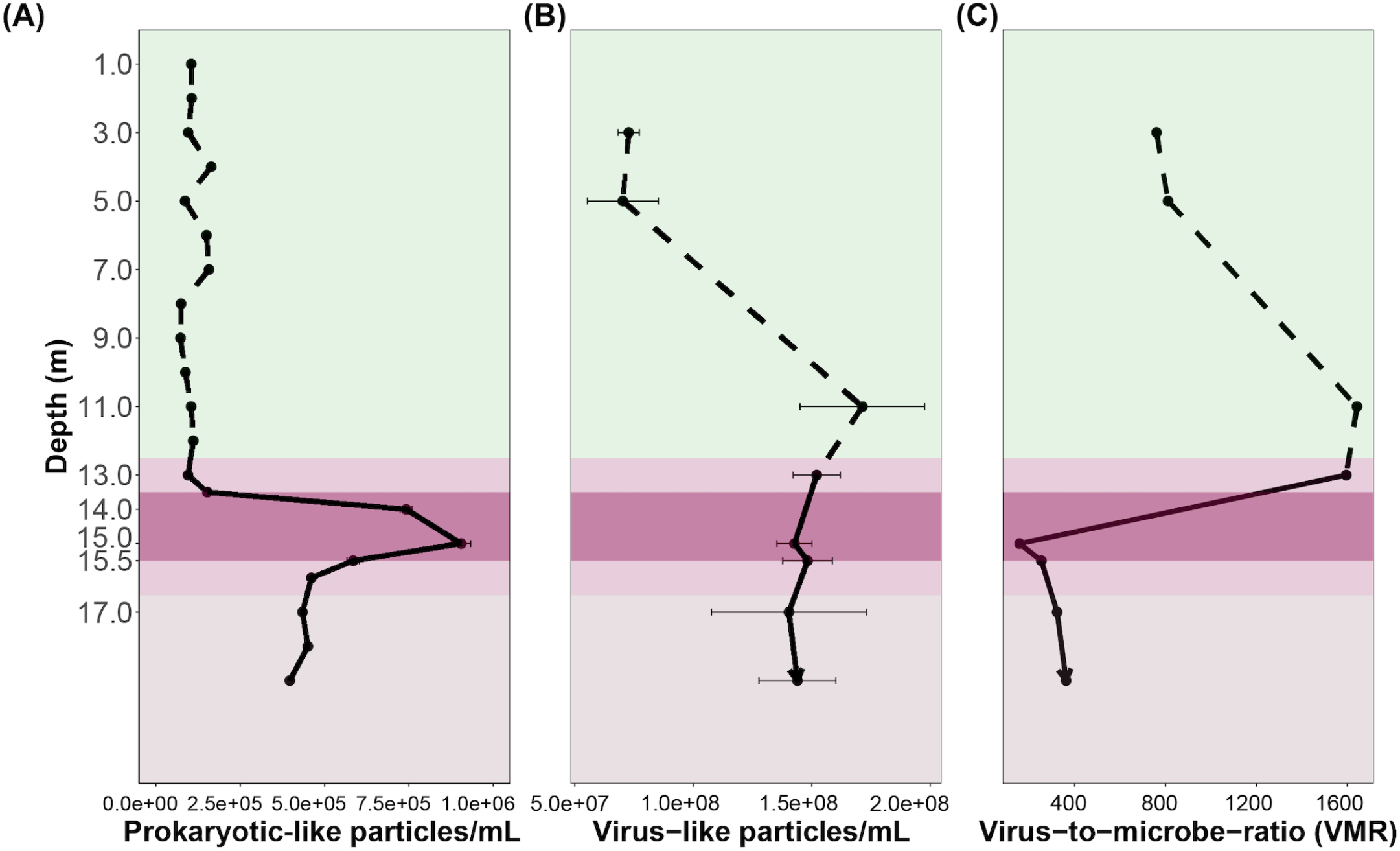
The abundances of (A) prokaryote-like particles (PLP) and (B) virus-like particles (VLP) from flow cytometry analyses and (C) virus-to-microbe ratio, as VLP/PLP.

Significant positive correlations were observed between PLP concentrations and other indicators of biomass: turbidity (R = 0.95; p-value = 0.00003), Chl a (R = 0.76; p-value = 0.01), POC R = 0.92; p-value = 0.0001), PON (R = 0.93; p-value = 0.0001), and POS R = 0.76; p-value = 0.010) (Fig. S4). Of those, only the VLP concentrations were correlated with Chl a (R = 0.77; p-value = 0.02, (Fig. S5). Significant negative correlations were observed between VLPs and oxygen (R = −0.82, p-value = 0.01) and light R = −0.90, 0.002). Significant positive correlations were observed between VLP and depth (R = 0.78, p-value = 0.02), conductivity (R = 0.92, p-value = 0.001), and SP (R = 0.70; p-value = 0.05) (Fig. S5).

### Bacterial community composition through the mixolimnion, chemocline, and monimolimnion

Both relative (OTU count scaled by total OTUs) and absolute abundances (OTU count scaled by total FCM PLP counts (29)) were considered to assess abundances of microbial taxa through Lake Cadagno (Fig, 3A, B). Absolute abundances of Bacteria classified cells (‘Bacteria’ OTU count scaled by total FCM PLP counts (29)) were at a minimum in the oxic mixolimnion (day 1; 0-12.5 m; Figure 2B, SI Table 1). On average, the mixolimnion community primarily consisted of Actinobacteria (35.34% of total OTUs; 33,815 cells/ml), Bacteroidetes (21.47% of total OTUs; 20,541 cells/ml), Proteobacteria (20.45% of total OTUs; 19,563 cells/ml), and Cyanobacteria (5.4% of total OTUs; 5,225 cells/ml) phyla (Figure 2A,B, Fig. S6).

In the chemocline, the absolute abundance of Bacteria-classified cells peaked at 15 m (day 2; Figure 3A), which corresponded with peak turbidity, POC and PON (Figure 1). The major phyla represented at this depth were Proteobacteria (48.42% of total OTUs; 438,272 cells/ml) and Bacteroidetes (25.35% of total OTUs; 229,448 cells/ml). The genera that dominated these phyla at 15 m were purple sulfur bacteria (PSB) Chromatium (35.18% of total OTUs; 318,381 cells/ml). Chromatium (Proteobacteria) was significantly positively correlated with turbidity (R = 0.93; p-value = 7.3e-5) and total cell counts (R = 0.99; p-value = 8.2e-09) (Fig. S7 A, B). An unclassified Lentimi-crobiaceae OTU, the second most abundant OTU in the chemocline at 15 m, was significantly positively correlated with secondary production (R = 0.8; p-value = 0.005), turbidity (R = 0.91, p-value = 0.0002), PLPs (R = 0.85, p-value = 0.001), PON (R = 0.71, p-value = 0.02), POS (R = 0.92; p-value = 5e-04), yet, negatively correlated with light (R = −0.78, p-value = 0.007) and oxygen concentration (R = −0.87; p-value = 0.001; Fig. S7 C-I). Though Firmicutes was not a dominant phylum itself, an unclassified genus of the phylum, *Erysipelotrichaceae* (3.90% of total OTUs; 35,360 cells/ml), was among the dominant genera at 15 m (SI Table 1). Green sulfur bacteria (GSB), *Chlorobium* (2.29% of total OTUs, 20,808 cells/ml), was the fifth most abundant genera of chemocline. Previously observed phototrophic sulfur bacteria, *Thiodictyon* and *Lamprocystis* (13, 10), and sulfate-reducing *Desulfocapsa* and *Desulfobulbus* were present at less than 0.01% of total OTUs in the chemocline (SI Table 1).

**FIG 3.**
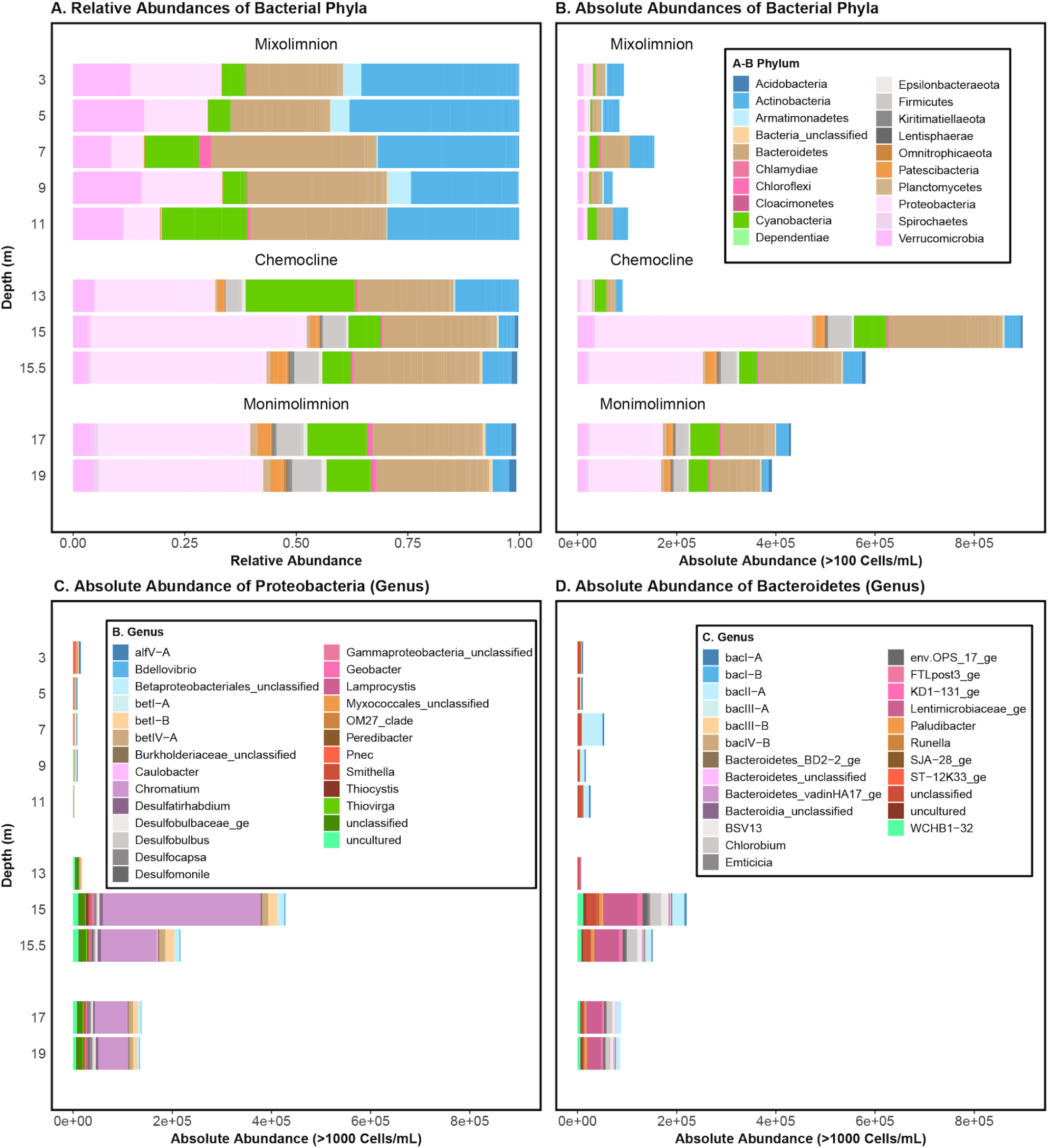
(A) Relative (OTU count scaled by total OTUs) and (B) absolute (OTU count scaled by total FCM PLP counts (29)) abundances of bacterial phyla at the sampled depths in Lake Cadagno, (C) genera in the Proteobacteria phylum, (D) genera in the Bacteroidetes phylum. To the right of each panel, the mixolimnion, chemocline transition zone, chemocline, and monimolimnion layers are indicated by light green, lilac, purple and light brown backgrounds, respectively and correspond with the depths reported on the left-most y-axis.

On average, the total PLP counts in the monimolimnion (17.0-19.0 m) were less than half that of the peak chemocline counts and more than twice the average mixolimnion counts. The average relative and absolute (Figure 3A,B) abundances of the dominant populations reflected those of the chemocline at 15 m, which included Proteobacteria (34.05% of total OTUs; 148,019 cells/ml), Bacteroidetes (24.53% of total OTUs; 106,635 cells/ml), Cyanobacteria (13.44% of total OTUs; 58,436 cells/ml), and Actinobacteria (5.8% of total OTUs; 25,301 cells/ml). In addition to bacteria, archaeal OTUs of the *Methanoregula* and *Woesearchaeia* genera were identified in the monimolimnion, but they were not further considered owing to their overall low abundances (<100 OTUs, SI Table 2).

### Oxygenic phototrophs in the mixolimnion, chemocline, and monimolimnion

Cyanobacteria was among the top phyla consistently observed at each depth in the water column, but at no depth did it dominate (Figure 3A). At all depths, *Oxyphotobacteria* and chloroplast-classified OTUs dominated the Cyanobacteria, representing 94-100% of the total Cyanobacteria-classified OTUs (Figure 4A). Of the low abundance cyanobacterial genera, Cyanobium was found throughout the water column, while *Pseudanabaena* and *Gastranaerophilales* were predominantly found at 15 m and deeper (Figure 4A-B). To better understand the relationship between the microbial guilds of Lake Cadagno, we sought more evidence regarding the origin of these putative chloroplast OTUs. The phylogeny of all Cyanobacteria-classified OTUs and their two nearest neighbours in SILVA (30) indicated that the chloroplast and *Oxyphotobacteria* OTUs clustered in clades with chloroplast 16S rRNA genes of cultured eukaryotic algae (Figure 4B): Chlorophyta (Otu00008, Otu000342, Otu00006, Otu00323), Ochrophyta (Otu00033), Streptophyta (Otu00208), Haptophyta (Otu00143), and uncultured-chloroplasts (Otu00042, Otu00051, Otu00052, Otu00055).

**FIG 4.**
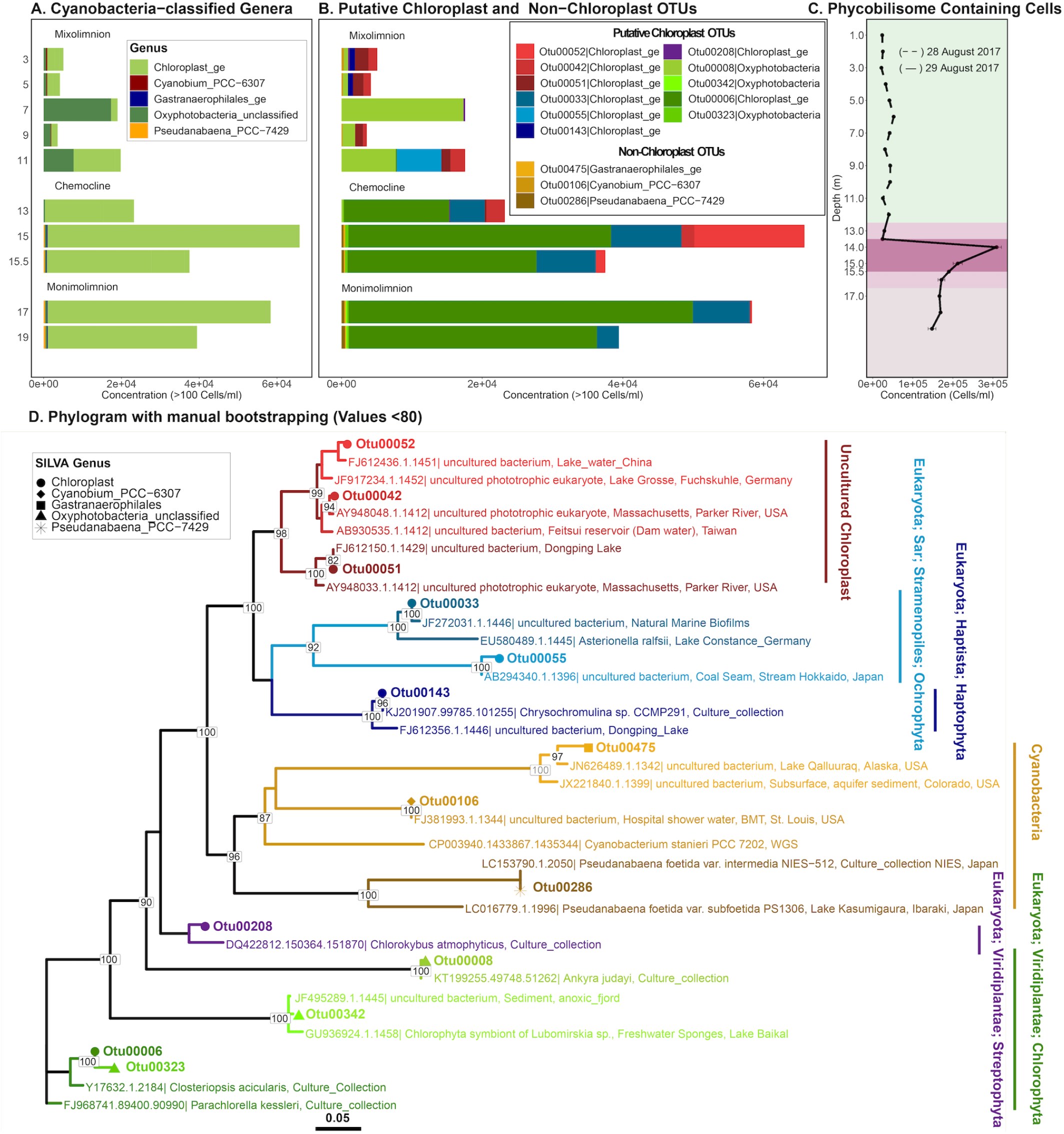
Abundances and phylogenies of Lake Cadagno phototrophs based on 16S rRNA gene amplicon sequencing and low-cytometry. Mixolimnion samples (0-11.0 m) were collected on day 1, chemocline and monimolimnion (12.0-19.0 m) samples were collected on day 2. Abundances in panels B-C are acquired by scaling 16S rRNA gene read counts by FCM cell counts. (A) Absolute concentrations (cells/ml) of OTUs >100 cells/ml assigned to the phylum Cyanobacteria, and of those, the (B) absolute concentrations (cells/ml) of OTUs classified as putative chloroplasts and non-chloroplast at the genus level. (C) Depth profile of phycobilins-containing cell counts based on low cytometry with a 640 nm laser. The mixolimnion, chemocline transition zone, chemocline, and monimolimnion layers are indicated by light green, lilac, purple and light brown backgrounds, respectively. (D) Phylogenetic tree of the representative 16S rRNA gene amplicon sequences of putative chloroplast and non-chloroplast OTUs in panel B, along with their two nearest neighbour sequences from the SILVA database. Clades are colour-coded consistent with the colouring of bar plots in prior panels.

In the lower mixolimnion between 7.0-11.0 m, where high oxygen, Chl a, and NPP persisted and peak Fv/Fm was observed (Figure 1D, F), Otu00008, which is most closely related to the chloroplast of cultured Chlorophyta, *Ankyra judayi*, dominated the subset of Cyanobacteria-classified OTUs where they represented up to 89.97% of the total cyanobacterial OTUs of mixolimnion (Figure 4B, D). At 11.0 m, the next most abundant OTUs were Otu00042 and Otu00055, which belonged to a clade of chloroplasts from uncultured organisms, representing 21.53% and 31.63% of the total cyanobacterial OTUs, respectively.

A primary chlorophyll peak was identified in the chemocline between 13.0-14.0 m, where FCM-based counts of phycobilin-containing cells (309,333 cells/ml; Figure 4C), phycocyanin pigments (26 μg/L day 2, Fig1D), and Chl a pigments (45 μg/L day 2) rose sharply (day 2; Figure 1D). While Cyanobacteria-classified OTUs represented nearly a quarter of all OTUs at the primary chemocline chlorophyll peak, they represented only 7.28% of the total OTUs at the secondary chemocline chlorophyll peak (15 m; SI Table 1). The most abundant chloroplast OTU throughout the chemocline, Otu00006 (Figure 4D), was most closely related to the chloroplasts of cultured Chlorophyta, *Parachlorella kessleri* and *Closteriopsis acicularis*, the latter a genus observed in meromictic Lake Tanganyika (31). The next most dominant chloroplast-like OTUs, Otu00033 and Otu00052, clustered with cultured Ochrophyta chloroplasts and uncultured chloroplasts, respectively (Figure 4D). In the monimolimnion, Otu00006 and Otu00033 dominated the Cyanobacteria-classified OTUs (Figure 4D), which were most closely related to Chloro-phyta and Ochrophyta chloroplasts, respectively.

### Prokaryotic genotypic and phenotypic diversity trends

Bacterial genotypic diversity (16S rRNA gene) and PLP phenotypic diversity (based on individual cell features distinguished by FCM (29)) both dropped at 7.0 m and rose in the lower mixolimnion where oxygenic primary production peaked (Figure 5A-B, Figure 1F). Overall, genotypic alpha diversity (Shannon) trends were near-uniform, except at the peak of turbidity (15.0 m) where PLP phenotypic alpha diversity peaked and moderate genotypic alpha diversity was observed (Figure 5A-B). Below the chemocline, consistently high genotypic and phenotypic alpha diversity was observed in the anoxic monimolimnion (Figure 5). In the Principal Coordinate Analysis (PCoA) ordinations of bacterial community dissimilarity (Bray Curtis), the first two components combined accounted for 77% and 46.2% of all observed variation in genotypic and phenotypic beta diversity, respectively (Figure 5 C-D). When both genotypic and phenotypic diversity was considered, the oxic mixolimnion and the co-clustering anoxic chemocline and monimolimnion samples separated along the first axis (Figure 5 C-D). In the phenotypic diversity ordination (Figure 5D), the mixolimnion samples separated along the second axis by whether they originated from the high-oxygen (1.0-7.0 m, light green) or the mid-oxygen (8.0-11.0 m, dark green) zones (Figure 5 C-D). As with phenotypic alpha diversity, oxygen and light were negatively correlated with phenotypic and genotypic beta diversity (Fig. S8).

**FIG 5.**
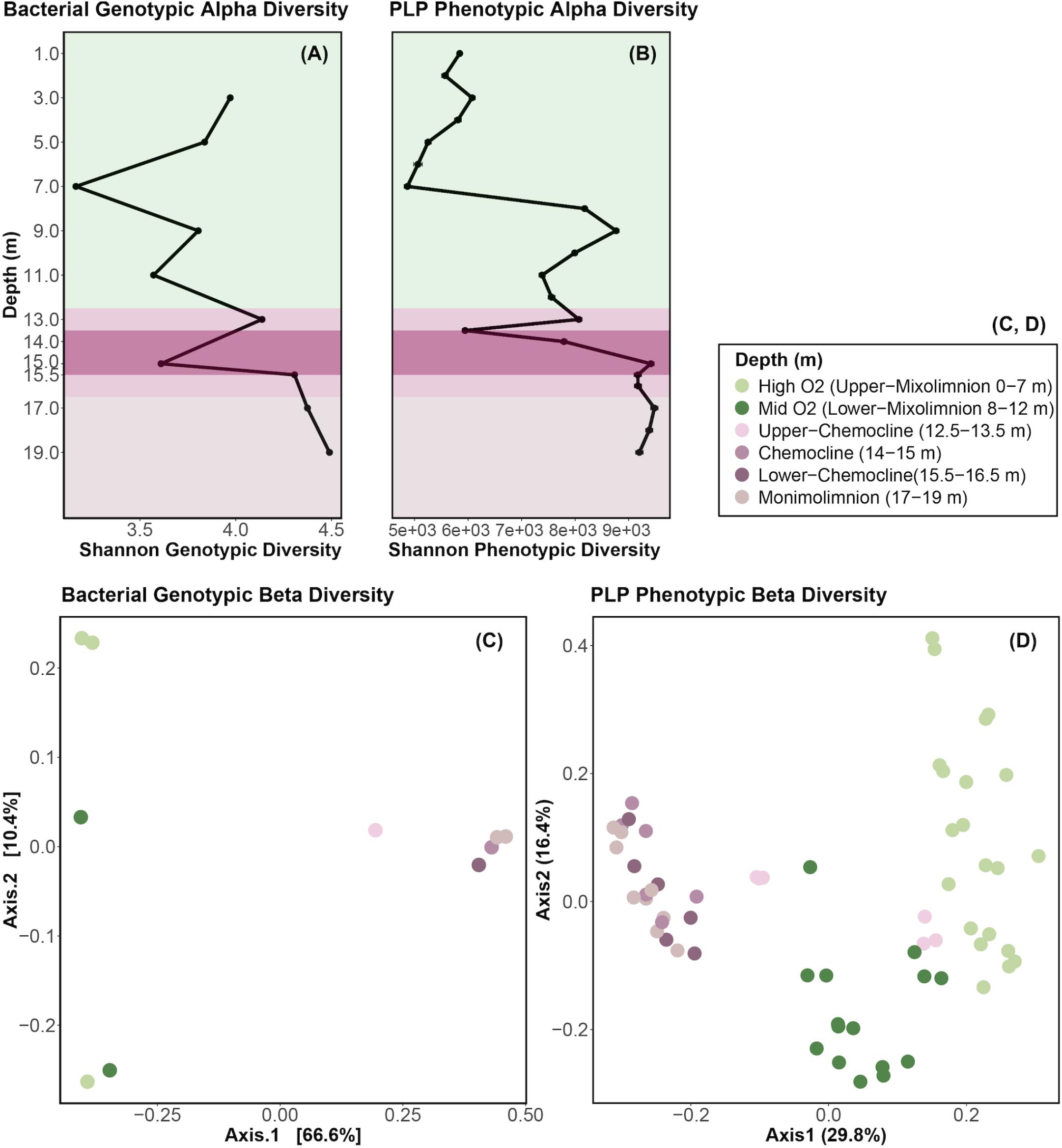
Trends in microbial community genotypic (16S rRNA gene amplicon sequencing, including bacterial and chloroplast-related OTUs) and phenotypic (PLP features based on flow-cytometry (29)) alpha and beta diversity in Lake Cadagno. (A) Variation in genotypic alpha diversity (Shannon) with depth. (B) Variation in PLP phenotypic alpha diversity (Shannon). Mixolimnion samples (0-11.0 m) were collected on day 1, chemocline and monimolimnion (12.0-19.0 m) samples were collected on day 2. Colours of data points represent depth strata sampled. (C) PCoA representing genotypic beta diversity dissimilarity (Bray Curtis) of microbial communities. (D) PCoA representing phenotypic beta diversity dissimilarity (Bray Curtis distance) of microbial communities.

## DISCUSSION

In this study, the investigation of viral, microbial and biogeochemical dynamics through Lake Cadagno’s water column improves our understanding of how biological and geochemical connections between the spatially segregated food webs in the Proterozoic ocean model (2) may have manifested.

### Eukaryotic algae associated with oxygenic phototrophy in the mixolimnion

With the major transition to an oxygenated ocean 800-700 Ma, the ever-reducing levels of toxic sulfide and relief of nitrogen limitations likely facilitated the evolution of modern eukaryote precursors, as proposed in the Proterozoic ocean model (2). In the oxic mixolimnion of Lake Cadagno, we observed low levels of sulfur and ammonia in combination with high abundances of eukaryotic algal chloroplast OTUs and photosynthetic activity, simultaneously with the low abundances of small (<40 μm) prokaryotic-like particles. These patterns reflect the conditions predicted during the early period of eukaryotic evolution in the Proterozoic ocean (Figure 6A).The chloroplast sequences identified in the mixolimnion were dominated by those of the fresh-water chlorophyte, *Ankyra judayi* (OTU00008; Figure 6B), which has not yet been described in Lake Cadagno. Likely due to its ability to endure high UV radiation, *Ankyra judayi* has been observed to outcompete other phytoplankton and grow to high densities in a high-alpine Andean lake (32), which may explain its presence in the sun-lit mixolimnion of high-alpine Lake Cadagno and may reflect traits of early photosynthesizing eukaryotes. Our findings are consistent with prior microscopy-based studies that have reported that the majority of the algal biomass was attributed to eukaryotic algae, such as *Echinocoleum* and *Cryptomonas* (Chlorophyta) and diatoms (16). These observations of the Lake Cadagno mixolimnion support the proposed evolution of eukaryotes in the oxic strata of the Proterozoic ocean, where, though spatially segregated, they co-occurred with abundant and productive chemocline and monimolimnion bacteria in a unified ecosystem.

**FIG 6.**
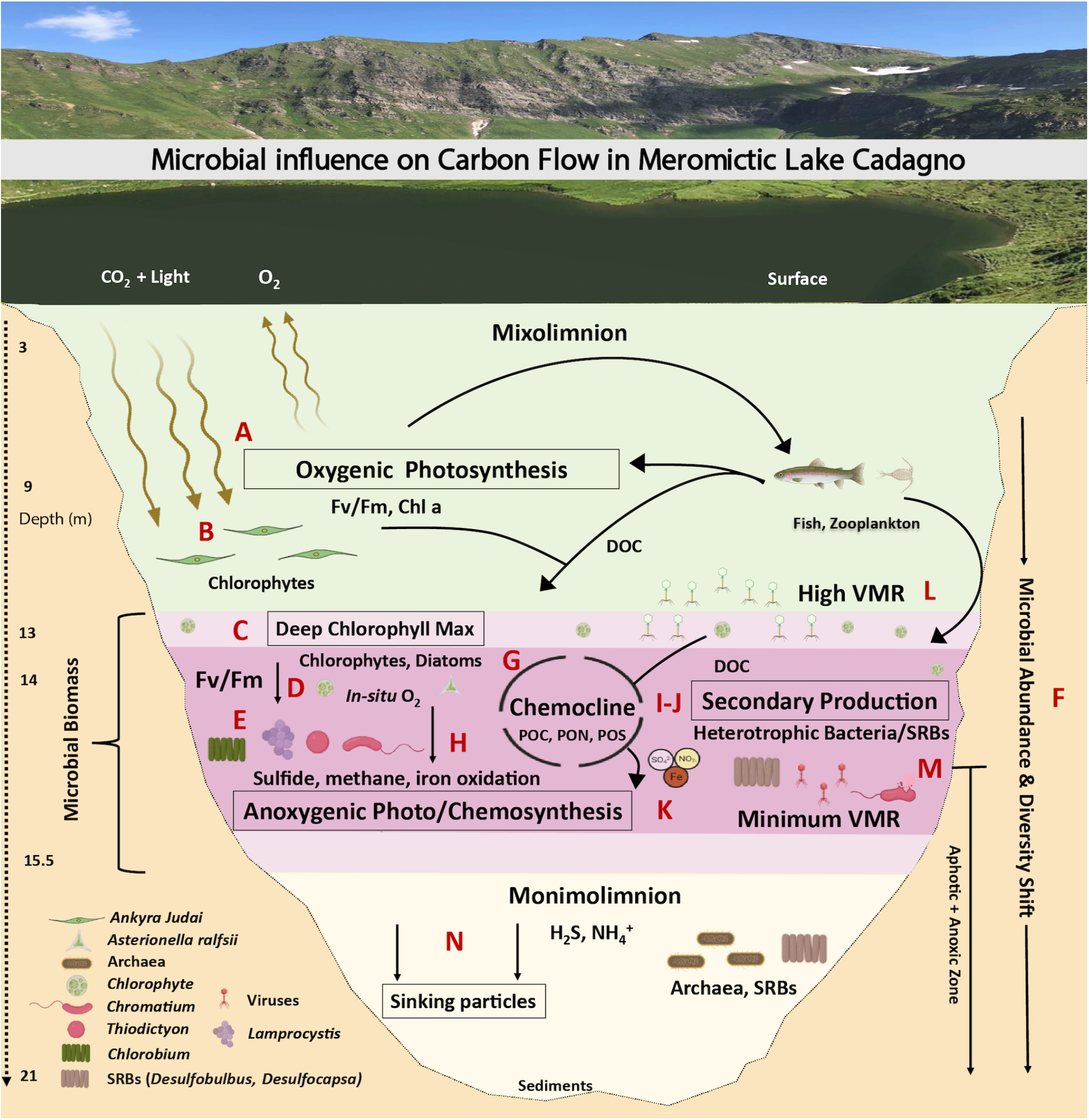
Microbial loop of meromictic Lake Cadagno. Microbiology icons created with BioRender.com.

### Coexistence of oxygenic and anoxygenic primary producers in the chemocline

The processes of the major microbial oxygenic and anoxygenic guilds have been reported to contribute almost equally to organic matter production in the Lake Cadagno chemocline (16), as they may have in ancient ocean chemoclines. Oxygenic (chlorophytes, diatoms, cyanobacteria) and anoxygenic phototrophs (PSB and GSB) coexisted in the Lake Cadagno chemocline (Figure 6C-E). Previous studies reported a cyanobacterial bloom (28 August, 12 September 2017) at the oxic-anoxic transition directly above the phototrophic sulfur bacteria (33, 34); however, cyanobacteria (Cyanobium, Gastranaerophilales, Pseudanabaena) were rare in this study. Furthermore, the shift of oxygenic to anoxygenic photosynthesis is coincident with a decreasing effciency of oxygenic photosynthetic microbes (Fv/Fm), increasing concentrations of hydrogen sulfide, and a shift in bacterial abundance and beta-diversity (Figure 6F). The nitrogen cycle likely mediated by N2 fixing cyanobacteria and purple and green sulfur bacteria has been proposed to provide inorganic nitrogen essential for ancient ocean photoautotrophy (2, 7). The peak in PON, and the rise in biologically available N (ammonia) at the peak turbidity, suggests active N-fixation was occurring in the Lake Cadagno chemocline. The major carbon assimilators of the chemocline, PSB (Chromatium, Thiodictyon) and GSB (Chlorobium), also perform nitrogen fixation (12, 35, 36, 7, 37) and are ancient clades thought to have been important sources of bioavailable nitrogen in Proterozoic oceans. Their diazotrophic metabolism likely provided the nitrogen needed for organic matter production (POC, PON, POS) in the Lake Cadagno chemocline (Figure 6G). Such relief from nitrogen limitation may have supported the persistence of PSB and GSB in the 1.6 Gyr old ocean basin of the permanently stratified Paleoproterozoic sea (38).

In addition to oxygenic and anoxygenic primary production, aerobic respiration has also been proposed to be coupled with biogeochemical processes of ancient ocean chemocline (2). However, evidence for which microbes are associated with such activity in the Proterozoic conceptual model is sparse. Though typically thought of for its role in photoautotrophy, Chromatium, the most abundant chemocline microbe in this study, is also known to perform light-independent chemolithoautotrophy and has been reported to contribute up to 40% of total dark carbon fixation through aerobic sulfide oxidation (39) (Figure 6L). In that study, aerobic respiration by Chromatium okenii co-occurred with in-situ production of oxygen by photosynthetic algae (39), methane oxidation by methanotrophs and iron oxidation, likely by *Chlorobium* and *Rhodobacter*, where iron oxidation accounted for 10% of total primary production in the chemocline (40, 41) (Figure 6H). Through their oxygen production, the oxygenic phototrophs (chlorophytes, diatoms, cyanobacteria) observed in the chemocline may play a critical role by enabling these oxidative metabolic processes central to Lake Cadagno’s biogeochemical cycling. These observations support the proposal of the Proterozoic ocean model of a mixed community of oxygenic and anoxygenic primary producers in the chemocline (2). Future work targeting the ecological and metabolic interactions between chlorophytes, diatoms, cyanobacteria and chemocline bacteria at the oxic-anoxic interface of Lake Cadagno will improve our understanding of the evolution of life in ancient oceans.

### Nutrient remineralization through secondary production contributes to chemocline biomass

While broadly recognized that the PSB and GSB that dominated the Lake Cadagno chemocline in this study are known to be largely responsible for anoxygenic primary production in this stratum (Figure 6E) (42, 10), we found that, as with primary production, secondary production rates were the highest in the chemocline, exceeding secondary production in the mixolimnion and monimolimnion by 42.4- and 1.9-fold, respectively (Figure 6I). This peak in secondary production in the zone of high biomass suggests that remineralization is important for supplying nutrients to the active chemocline microbial assemblage. By pairing these secondary production rates with a description of heterotrophic populations present, we shed light on the clades that may be involved in nutrient remineralization in the chemocline. Lentimicrobium, the second most abundant genus in this report, has been implicated in fermentation under limited light and oxygen in previous studies (43). Sulfate-reducing bacteria (SRBs),(Figure 6J) such as the observed Desulfocapsa and Desulfobulbus, are known to form aggregates with phototrophic PSB (e.g., Thiodictyon syntrophicum, Lamprocystis purpurea) and both co-occurred in the chemocline (44, 45). Through secondary production, heterotrophic microbes are likely involved in the decomposition of particulate organic matter (POC, PON, POS) and the supply of inorganic nutrients (iron, sulfate, and nitrogen) necessary to carry out primary production by and sustaining populations of oxygenic and anoxygenic photoautotrophs in the chemocline (Figure 6K). These autotrophic and heterotrophic relations combined with the secondary productivity rates provide the evidence for which microbes may have been involved in the recycling of organic matter proposed in the Proterozoic ocean model (2).

### Potential for role of viruses in modulating microbial activity and nutrient cycling

Viruses serve as top-down controls on host populations and reprogram host metabolism during infection (19, 46, 20). Evidence for viral activity has been observed in ancient cyanobacterial mat analogues, including their potential for resource scavenging through the degradation of host phycobilisomes, a pigment central to photosynthesis (47). In Lake Cadagno, VMR peaked in the lower mixolimnion and mixolimnion-chemocline transition where high VLP and chlorophyll concentrations and high photosynthetic activity were observed together with low bacterial abundances. This colocalization of high viral loads and highly active phytoplankton populations could be explained by ongoing lytic viral infections of phytoplankton near the DCM (Figure 6L). Such a scenario was observed in the phytoplankton Ostreococcus virus-host system, where host densities were maintained concurrently with continual lysis due to phase switching between resistant and susceptible host, a strategy supported by mathematical modelling (48). If this scenario is occurring, the sustained lysis of phytoplankton has the potential to release DOC in a viral shunt near the mixolimnion-chemocline transition. Such a DOC peak was observed slightly below this depth in Lake Cadagno. However, given that VMR is a feature that emerges from a number of underlying virus-microbe interactions, viral life-history traits, and environmental co-factors (49), the prediction of sustained phytoplankton viral predation at the mixolimnion-chemocline transition requires empirical confirmation.

Previous studies have identified genomic evidence for viral infection of the presumed major carbon assimilators in the chemocline, Thiodictyon syntrophicum, and Chromatium okenii (10). Genomes of these organisms sequenced from Lake Cadagno contain CRISPR elements (35, 36), which derive from a type of acquired immunity against invading viral and plasmid DNA. This suggests viruses may play a role in the modulation of carbon, and sulfur cycles, as has been proposed in hydrothermal vent (50) and wetland (51) microbial communities, where viruses have been found to carry genes central to methanogenesis (mcrA) and sulfur reduction (dsrA, dsrD). Active viral lysis also provides a new possible mechanism to explain the previously observed low abundance of Chromatium okenii, which has previously been attributed to microbial predation (15). Despite genomic evidence suggesting infection of these populations, VLP counts did not rise with PLP counts in the chemocline. Yet, our observation of minimum VMR (Figure 6M) and peak microbial abundances in the Lake Cadagno chemocline supported an emerging trend in stratified meromictic lakes, as the same observation was made in meromictic Ace Lake (Vestfold Hills of East Antarctica) (23). Such coincident low VMR and high microbial abundance have been recognized as a hallmark of the power-law relationship proposed to describe the VMR dynamics in aquatic systems (49). We propose viruses as an important component of the Proterozoic ocean microbial community. Population-specific studies of viral infection are needed in Lake Cadagno to understand the impact of viruses on the evolution and function of microbes that underlie its biogeochemistry and to shed light on theroles viruses may have played in the ancient ocean.

## CONCLUSION

This work highlights how biogeochemical exchange within microbial guilds may be driving biomass accumulation and support the greater food web of the permanently stratified Lake Cadagno. Ultimately, the organic matter generated through eukaryotic and bacterial activity and viral remineralization in the chemocline can be made available to zooplankton and fish, whose biomass eventually sink to the sediments contributing to the organic matter burial (Fig. 6N). The observed trends suggest a high degree of interconnectedness and total ecological importance of microbial guilds in stratified ancient oceans.

This study inspires future research directions towards a better ecological and biogeochemical understanding of Lake Cadagno and similar meromictic ancient ocean analogues. Our community composition and secondary production data suggest that the ecological role and metabolic potential of heterotrophic bacteria that recycle chemocline biomass in the presence of sulfur and ammonia, such as the newly identified and abundant Lentimicrobium, are intriguing unknowns. Furthermore, eukaryotic phytoplankton are known to contribute to the deep chlorophyll max of Lake Cadagno’s chemocline for two decades (16), insights into the cellular machinery that allows them to photosynthesize under limited light and oxygen remain unknown. Further, while primary producers (phytoplankton, phototrophic sulfur bacteria) and heterotrophic bacteria are central to the realized function of the lake biogeochemistry and ecology, viruses may shuffle the repertoire of genes controlling both microbial guilds, hence modulating biogeochemical (carbon, nitrogen and sulfur) cycles of the lake. Future work will build from our empirical observations to better understand the role that microbes of the chemocline (oxic-anoxic boundary) may have played in the Proterozoic transition from primarily anoxygenic to oxygenic metabolisms that dominate nutrient biogeochemical cycling in modern oceans. (52, 2, 53)

## MATERIALS AND METHODS

### Water-sample collection and Physicochemical profiling

This study was conducted in Lake Cadagno (21 m deep), a high alpine meromictic lake situated at 1921 m above sea level in the Southern Alps of Switzerland (54). Water sampling and characterization of physical (turbidity, temperature, oxygen, conductivity), biological (Chl a, phyco-cyanin) and chemical (H_2_S, NH_4_^+^) parameters of the water column were performed as previously described (11). Water was sampled on the 28 (Day 1) and 29 (Day 2) August 2017 using a double diaphragm teflon pump (Almatech PSG Germany Gmbh) connected to acid-washed low-density polyethylene (LDPE) line deployed from a platform that was anchored above the deepest part of the Lake Cadagno (46.55087° N, 8.71152° E). The oxic mixolimnion (0-10 m) was sampled on 28 August (Day 1), and the chemocline (11-16 m) and monimolimnion (17-19 m) were sampled on 29 August 2017 (Day 2).

Vertical lake profiles were determined using an autonomous conductivity temperature depth sensor (CTD; Ocean Seven 316 Plus CTD; IDRONAUT, S.R.L.). The CTD recorded pressure (dbar), temperature (oC), and conductivity (mS/cm); dissolved oxygen (mg/L) was measured with a pressure-compensated polarographic sensor. The error in oxygen profiles for day two was manually adjusted by referring to day 1 measurements which indicated approximately zero oxygen within and below chemocline. CTD profiles from the whole water column (0-20 m, Day 1 and 2) were used to identify the distribution of the mixolimnion, chemocline and monimolimnion and guided water sampling strategy. The CTD was equipped with an LI-192 Underwater Quantum Sensor (Li-Cor Biosciences; NE, USA) that continuously recorded photosynthetically active radiation (PAR-w/m2;) and a TriLux multi-parameter algae sensor (Chelsea Technologies Ltd; Surrey, UK) that measured in-vivo Chl a (Chl a, μg/L), an indicator of phytoplankton (including eukaryotic autotrophs and cyanobacteria) biomass (55), and phycocyanin (μg/L), a pigment characteristic of cyanobacteria (56).

### Chemical parameters

#### Dissolved compounds

Samples for the analysis of dissolved compounds (DOC) were filtered through an Acropack filter cartridge (PALL) with a 0.8 μm pre-filter and 0.2 μm final filter. Filters were collected in 30 mL acid-washed and pyrolyzed pyrex tubes filled to the top and supplemented with 100 μL HCl 2M. Tubes were stored in the dark at 4° C until processed (November 2017; University of the Geneva) using a Shimadzu TOC-LCPH analysis system. Blanks consisting of Milli-Q water were made and calibration was done using the standard from 0.2 to 5 ppm. Dissolved (NH_4_^+^)) and (H_2_S) were analyzed onsite at the Alpine Biology Centre (CBA; Piora, Switzerland) with freshly collected water samples using a UV-visible spectrophotometer (DR 3800, HACH) (57).

#### Particulate compounds

Particulate organic carbon (POC), nitrogen (PON) and sulfur (POS) were collected by filtering 150 to 500 mL of lake water (volume necessary to begin to clog the filter) on pre-combusted 47 mm GF/F filters (5h at 550 °C, cat. WH1825-047; Whatman, Wisconsin, USA), which were then acidified (500 μL HCL 1 M, added twice at 1 h interval) and dried in an oven (65 °C) overnight in acid-washed pyrex petri dishes. Samples were sealed using parafilm and aluminium foil until analysis (November 2017, University of Geneva) using an Elemental Analyzer (2400 series II CHNS/O Elemental Analysis, PerkinElmer). Procedural blanks using MilliQ water were performed (24-25 November 2017) in duplicate to measure the background signal, which was subtracted to sample analysis. POC, PON, POS were expressed in mg/L (ppm).

### Biological parameters

#### Primary production

Primary productivity was determined using incubation with NaH14CO3 (Perkin Elmer cat. NEC086H005MC; Waltham, MA, USA; 5 mCi, 1 mL in glass ampoule) that was freshly diluted into 4 mL MilliQ water at pH 9.6 (adjusted using NaOH, Sigma) and added at a dose of 1 mCi/L of lake water, as described (58). Incubation was carried out along an incubation chain deployed at six depths for 23 h on 28-29 August 2017. Each depth consisted of a metallic ring fixed to a rope at the desired depth (3, 5, 7, 9, 11 and 13 m). The ring was surrounded by six arms on a horizontal plane, each of which ended with a bottle holder. 75 mL acid-washed glass bottles were filled to the top with lake water (ca. 100 mL) at the corresponding depth and spiked with 14C prior to being deployed on each level of the incubation chain. At each depth, three transparent bottles were incubated for activity measurements at in situ natural light intensities, and three amber bottles were incubated for dark community respiration measurements. At the end of the incubation, each bottle was filtered onto 47 mm GF/F filters (cat. WH1825-047, Whatman, Wisconsin, USA) then each stored in a plastic petri dish. Petri dishes were placed in a sealed box containing calcium hydroxide (slaked lime) powder. After adding HCL (1M, 500 μL) onto filters, inorganic carbon degassed overnight in a sealed box. Filters were then collected into 20 ml scintillation vials, supplemented with 10 mL of Ultima Gold AB cocktail, and shaken manually. Primary productivity was expressed in μmol of net carbon fixation per h either per L of lake water or per Chl a, considering a dissolved inorganic carbon concentration of 1.8 mM C (14).

#### Maximum quantum yield

The maximum quantum yield (Fv/Fm) informs the photo-synthetic health and biological activity of algae based on photo-physiological characteristics of the chlorophyll photosystem II. Maximum quantum yield was determined using a Fast Repetition Rate Fluorometer (FRRF, FastOcean PTX coupled to a FastAct base unit; Chelsea Technologies). Water for maximum quantum yield measurements was pre-concentrated 10-fold by gently resuspending the cells as the sample passed through a 0.22 μm 47 mm polycarbonate filter (cat. GVWP04700, Millipore; Darmstadt, Germany) using a hand pump (pressure below 15 mbar) and a 47 mm filtration unit (cat. 300-4000; Nalgene). This step was done to ensure high enough method sensitivity and has been shown to alter neither Chl a nor cell integrity of samples from Lake Geneva (59). The FRRF was used in single turnover mode to record, in a 45 min dark-adapted natural sample, the basal (Fb) and the maximal (Fm) fluorescence following exposure to intense light. The sample was then gently filtered on a 0.22 μm 47 mm filter (cat. GVWP04700, Millipore; Darmstadt, Germany) to record the residual fluorescence (Fr). For each sample, the Fb-Fr represented the initial dark-adapted fluorescence F0 associated with intact cells. The Fv/Fm was then calculated using the ratio: (Fm- F0)/Fm.

#### Flow-cytometry (FCM)

A BD Accuri C6 cytometer (Becton Dickinson, San José CA, USA) with two lasers (blue: 488nm, red: 640 nm) and four fluorescence detectors (lasers 488 nm: FL1 = 533/30, FL2 = 585/40, FL3 = 670; laser 640 nm: FL4 = 675/25) was used to estimate total cell counts from fresh unpreserved samples collected on-site. FL4 detectors were specifically used to detect emission from phycobilin, a pigment proxy for cyanobacteria. Cells were stained with SYBR green I (cat. S7563, Molecular Probes; Eugene, OR) with a ratio of 1:10,000 (vol/vol), following incubation for 13 minutes at 37°C in the dark (9). Histogram of event counts versus green fluorescence (FL1 > 1,100) allowed quantification of total cells.

VLPs were counted using flow-cytometry after filtration of lake water through 55 μm mesh and 0.22 μm filters to remove large particles and cells. In triplicate, 1 ml sample from the filtrate was fixed by adding 10 μL of glutaraldehyde (25% stock solution) in sample cryovials. The samples were fixed for 15 minutes at room temperature and subsequently stored in liquid nitrogen for shipping, then stored at −80°C until processed in February 2018. Virus-like particle counts were obtained using a FACSCalibur flow cytometer (Becton Dickinson Biosciences; Grenoble, France). Samples were thawed at 37°C then diluted in autoclaved and 0.02 μm filter-sterilized Tris-EDTA (0.1 mM Tris-HCL and 1 mM EDTA, pH 8). Samples were then stained for 10 minutes at 75°C using SYBR Green I (Molecular Probes) at a 1:10,000 final concentration (60).

#### Phenotypic alpha diversity of PLP (Prokaryotic-like-particles)

Raw flow-cytometry files were used to estimate phenotypic traits (morphology and nucleic acid content) using the Phenoflow package (29) in R (v4.1.0) (61) and R-Studio (v1.4.1106) (62). Briefly, this approach estimates kernel densities on multiple bivariate single-cell parameter combinations (e.g., fluorescence and scatter intensity) and concatenates these into a feature vector, which can be thought of as a phenotypic fingerprint (29). This fingerprint represents community structure in terms of the phenotypic attributes of the cells that comprise the community—essentially, any feature that influences the way the laser interacts with passing particles, such as morphology, size, and nucleic acid content, is captured in the phenotypic fingerprint. Using this approach, bacterial phenotypic diversity metrics have been shown to be highly correlated with community taxonomic diversity (29). To calculate phenotypic alpha diversity, phenoflow calculates Hill number diversity indices (Dq) from order 0-2, where 0 represents the richness and q>1 represents richness and evenness, also referred to as D0 (richness), D1 (Shannon), and D2 (inverse Simpson) (63). Phenotypic beta diversity was estimated using PCoA of phenotypic fingerprints.

Fluorescence detectors (lasers 488 nm: FL1-H = 533/30, FL3-H = 670) were used to target PLP diversity. Polygon gating was applied to filter out the noise observed in control samples (Fig. S1). Clockwise, the polygon gate coordinates (x,y) were: (7, 6), (15, 6), (15, 17), and (7,10). Kernel density estimates were calculated by applying a binning grid (128 × 128) to FSC-H (Forward scatter height), SSC-H (Side scatter height), FL1-H and FL3-H. The obtained kernel densities values were assigned into one-dimensional vectors, termed phenotypic fingerprints.

#### Secondary production

Heterotrophic bacterial production, also termed secondary production, was estimated by the micro-centrifuge 3H-Leucine bacterial production method (64). A working solution of 5 μCi mL-1 leucine was made from a manufacturer stock solution of radioactive leucine, L-[3,4,5-3H(N)] (cat. NET460A005MC, PerkinElmer, MA, USA). To reduce the potential for autodegradation of the leucine radiolabel, stock 3H-Leucine material was stored in the dark at 2 to 4° C (never frozen) and used for the incorporation experiment within 5 days. Two replicates and one trichloroacetic acid (TCA)-killed control (5% [vol/vol] final concentration; cat. 91228, Sigma) were used for each depth. Briefly, 1.5-mL of lake water samples were incubated with a mixture of radioactive and nonradioactive leucine at final concentrations of 20 nM. Using the [3H]-Leucine working solution, enough “hot” leucine was added to each tube to create a sufficient signal (50% or 85% of hot leucine were used in the mixture for samples collected above and below 13m depths, respectively). Incubations were conducted for 2 hr in a dark incubator at in situ temperatures that corresponded to their sample depths. Saturation of leucine incorporation over this period was tested with 20, 30 and 40 nM of total leucine. At the end of the incubation, 200 μL of 50% TCA is added to all but the control tubes to terminate leucine incorporation. To facilitate the precipitation of proteins, bovine serum albumin (BSA; Sigma, 100 mgL-1 final concentration) was added, then samples were centrifuged at 16,000 g for 10 min (65). The supernatant was discarded and the resultant precipitated proteins were washed with 1.5 mL of 5% TCA by vigorous vortexing and again centrifuged (16,000 g for 10 min). The supernatant was discarded. Subsequently, 1.5 mL of UltimaGoldTM uLLt (Part number: 6013681, PerkinElmer, MA, USA) was added to each vial, mixed, and allowed to sit for >24 h at at room temperature before the radioactivity was determined using a liquid scintillation counter (Beckman LS6500). To calculate biomass production, the following conversion factors were applied: molecular weight of leucine of 131.2 g/mol, fraction of leucine per protein of 0.073, the ratio of cellular carbon to the protein of 0.86, and 2.22×106 DPM/μCi to convert disintegrations per minute (DPMs; the measure of radioactivity) to μCi. A factor of 1.55 kg C mol leucine-1 was used to convert the incorporation of leucine to carbon equivalents, assuming no isotope dilution (66).

#### Statistical Analyses

The magnitude and significance of relationships between physical and biological parameters were determined with linear regressions and visualized with scatter plots using R (v4.1.0) (61) in R-Studio (1.4.1106) (62) using ggplot2 (3.3.3) (67) and ggpubr (0.4.0) (68) packages (69). R > 0.70 was considered a strong positive correlation, and a p-value < 0.05 was considered statistically significant.

### Illumina MiSeq 16S rRNA gene amplicon sequence analyses

#### 16S rRNA gene amplicon sequencing

Water samples collected from 3.0, 5.0, 7.0, 9.0, 11.0, 13.0, 15.0, 15.5, 17.0 and 19.0 m (20 L/depth) were 55 μm pre-filtered to remove large particles and then 0.22 μm filtered (cat. GPWP14250 142 mm Express Plus Filter, Millipore; Darmstadt, Germany) using a peristaltic pump. Filters were flash-frozen in liquid nitrogen and stored at (−80°C) until extraction of DNA using the DNeasy Blood and Tissue Kit (cat. 69504 QIAGEN, Germantown, MD, USA) combined with QIAshredder (cat. 79654, QIAGEN, CA, USA), as described (70). The first and second elutions of DNA (50 μl each) were stored (4o C) until sequencing. Dual indexed primers were used and the V4 region of the 16S rRNA gene amplicon was targeted and amplified, as described (71). Sequencing libraries were prepared using the Illumina Nextera Flex kit (Illumina; CA, USA) and sequenced using the MiSeq platform (500 cycles, Illumina) at the University of Michigan Microbiome Core facility. The sequenced reads were quality controlled, assembled, trimmed, aligned, and clustered at 97% identity threshold into a representative operational taxonomic unit (OTU) using mothur (72). The OTUs were taxonomically classified using SILVA (73) and TaxAss freshwater(74) 16S rRNA gene databases.

#### Genotypic alpha and beta diversity and Phylogenetic Tree

Downstream analysis was carried out by the Phyloseq package (75) using R (v4.1.0) (61) in R-studio (1.4.1106) (62) to analyze the alpha and beta genotypic diversity of bacterial communities. Taxa not observed in 20% of samples at least three times were removed. The Alignment, Classification, and Tree (ACT) function of SILVA was used to generate the phylogenetic tree (76). Representative sequences of chloroplast OTUs were aligned to SILVA sequences (SINA 1.2.11) Two neighbours per query sequence (95% minimum sequence identity) were added to the tree with the chloroplast OTU sequences. Advance variability profile was selected as ‘auto’ and sequences below 90% identity were rejected. The phylogenetic tree of the OTUs was bootstrapped with RAxML (77) with 20 maximum likelihood (ML) and 100 bootstrapped searches. The tree was annotated using the R package ggtree (2.2.4) (78) and Adobe Illustrator (25.2.1).

#### Absolute Abundances

The absolute quantifications of taxa were determined as described (79). Briefly, the relative abundance of bacterial and archaeal OTUs of a given taxon (taxon read count/total read count in sample) was multiplied by the PLP concentrations (cells/ml) for that sample, as determined by flow cytometry (FCM). Thereon, archaeal abundance was deducted from the total prokaryotic population to get an absolute abundance of the bacterial population. Chloroplast OTUs were assigned to the phylum Cyanobacteria by SILVA, owing to their cyanobacterial origins ((80).

### Data Availability

Raw sequences are deposited in the NCBI Short Read Archive (SRA) and available through the BioProject PRJNA717659. The code used for running the mothur analysis, statistical analyses, and figure generation are available at the project GitHub repository: https://github.com/DuhaimeLab/Lake_Cadagno_microbial_loop_Saini_et_al_2021.

## SUPPLEMENTAL MATERIAL

**Table S1.** The relative and absolute abundance of bacterial populations, including chloroplasts.

**Table S2.** The relative and absolute abundances of archaeal populations.

**Figure S1.** Raw flow-cytometry files visualized with BD6 flow-cytometer interface (A-B) and R-studio Phenoflow package (A-D). The red outline represents the gating strategy for cell counting and phenotypic diversity analysis. FCM events representing prokaryotic-like particles in a Lake Cadagno (1 m depth) targeted using (A) automatic gating via a four-quadrant layout and (B, C) manual gating. (D) FCM events in a 0.2 μm filtered, cell-free MilliQ water (control).

**Figure S2**. Experiment to monitor the health of photosynthetic cells. Basal fluorescence (Fb) and maximum fluorescence (Fm) were obtained to calculate Fv/Fm values.

**Figure S3**. Line plots, R-coefficients, and p-values of linear regressions performed between secondary production, photosynthetic pigments, Prokaryotic-like-particles (PLPs), and physicochemical parameters, including turbidity, oxygen and light. Lake depth (m) is indicated by point colors. (A) Secondary production rates vs Chl a concentrations (B) Secondary production vs phycocyanin concentrations (C) secondary production rates vs turbidity (D) Secondary production vs Oxygen concentrations (E) secondary production rates vs light levels (F) secondary production rates vs PLP concentrations (G-J), Chl a and phycocyanin concentrations vs oxygen concentration and light levels. Grey zones represent 95% confidence intervals.

**Figure S4**. Line plots, R-coefficient, and p-values of linear regression performed between Prokaryotic-like-particles (PLPs) and biomass indicating factors. Lake depth in meters (m) is indicated by colors. (A) PLPs vs turbidity (B) PLP concentrations vs Chl a concentrations (C) PLP concentrations vs particulate organic carbon (POC) concentrations. (D) PLP concentrations vs particulate organic sulfur (POS) concentrations. (E) PLP concentrations vs particulate organic nitrogen (PON) concentrations. Grey zones represent 95% confidence intervals.

**Figure S5**. Line plots, R-coeffcients, and p-values of linear regressions performed between Virus-like-particles (PLPs) and physicochemical parameters. Lake depth (m) is indicated by colors. (A) VLP concentrations vs oxygen concentrations (B) VLP concentrations vs light levels (C) VLPs vs depth. (D) VLP concentrations vs conductivity. (E) VLP concentrations vs secondary production rates. Grey zones represent 95% confidence intervals.

**Figure S6**. Absolute abundances of microbial communities at the genus level through the vertical water column of Lake Cadagno.

**Figure S7**. Line plots, R-coefficients, and p-values of linear regressions performed between microbial abundances and physicochemical parameters. Lake depth (m) is indicated by point colors. (A-B) Chromatium abundances vs PLP concentrations and turbidity. (C-I) Lentimicrobium abundances vs secondary production rates, turbidity, PLP concentrations, light levels, and PON, POS, and oxygen concentrations. Grey zones represent 95% confidence intervals.

**Figure S8**. Line plots, R-coefficients, and p-values of linear regressions between microbial alpha and beta diversity and physicochemical parameters. Lake depth (m) is indicated by point colors. (A-B) PLP phenotype-based alpha diversity (Shannon) vs oxygen concentrations and light levels. (C-D) 16S rRNA gene-based beta diversity vs oxygen and light levels. (E, F) PLP phenotype-based beta diversity vs oxygen concentrations and light levels. (G-H) Phenotypic vs genotypic prokaryotic alpha and beta diversity. Grey zones represent 95% confidence intervals.

## ACKNOWLEDGMENTS

The project was funded by Swiss National Science Foundation grant no. PP00P2-138955 and through financial support to J.S.S. from the Swiss Confederation and Ernst and Lucy Schmidheiny Foundation that graciously enabled the development of this manuscript. The funders had no role in study design, data collection and interpretation, or the decision to submit the work for publication.

We thank the Duhaime Lab at the University of Michigan for supporting a learning environment to carry out the sequencing, flow cytometry, and ecological analyses and for productive feedback during the manuscript development. Many thanks to Dr. Sandro Peduzzi and the Alpine Biology Center Foundation (Switzerland) for providing accommodations during Lake Cadagno sampling and for offering comments on the manuscript. The University of Michigan Microbiome Core facility aided in the generation of Lake Cadagno 16S rRNA gene data.

